# Prolonged inhibitory effect of repeated period tibial nerve stimulation on the micturition reflex in the rat

**DOI:** 10.1101/2020.07.07.191213

**Authors:** Jiawen Zeng, Shaohua Zeng, Chonghe Jiang, Sivert Lindström

**Author notes:** These authors were co-authors (Jiawen Zeng, Shaohua Zeng). Corresponding author: Professor Chonghe Jiang.

## Abstract

**Background:** The aims of this study was determine if stimulation of tibial nerve afferents could induce a prolonged modulation of the micturition reflex in the rats.

**Methods:** Fifteen female Sprague Dawley rats (250-350 g) were fully decorticated and paralysed for the study. Tibial nerve stimulation (TNS) was delivered by inserting two pairs of needle electrodes close to the nerves at the level of the medial malleolus. Constant flow cystometries (0.07 ml/min) at about 10 min interval were performed and the micturition threshold volume (MTV) was recorded and used as the dependent variable. After 4 – 5 stable control recordings, the tibial nerves of both sides were stimulated continuously for 5 min at 10 Hz, 3 times threshold for α-motor axons. Six times of same stimulation were applied repeatedly with an interval of 5 min between the stimulations. The mean MTV was compiled from several cystometries in each half hour before the TNS and during, after 6 periods TNS.

**Results:** During the experiment, all the animals survived in a good condition with reasonably stable micturition reflexes, a significant increase in MTV was revealed after TNS. The best effect (mean 178%) occurred during the first 30 min after 6 periods of stimulation. This clear threshold increase remained for at least 5 h.

**Conclusions:** A prolonged increase in MTV was demonstrated by a short periods of TNS repeatedly. This post stimulation modulatory effects of micturition reflex would provide a theoretical explanation for the clinical beneficial effect of TNS in patients with overactive bladder (OAB).

## Introduction

Tibial Nerve Stimulation (TNS) is defined as a minimally invasive procedure by posterior tibial nerve stimulation percutaneously and delivering electrical stimulation to sacral nerve plexus. Activating this afferent could induce sympathetic inhibitory pathway as well as central inhibitory connection to the parasympathetic preganglionic neurons to suppress the bladder overactivity [1-3].

Clinical studies showed that TNS induced a prolonged poststimulation inhibitory effect on bladder activity lasting for days or weeks [4-6]. Therefore, this procedure has been listed as optional therapy in the AUA treatment guidelines as the third-line treatment for overactive bladder (OAB) [7]. Despide the beneficial effect of this procedure has been reported clinically, the underlying mechanism of action remains to be clarified.^4^ Experimental studies on urinary bladder physiology in cats and rats demonstrated that TNS could induce an immediate as well as prolonged bladder inhibitory effect manifested as increase in bladder capacity and decrease in micturition frequency. However, the data showed that the poststimulation effect only within a short time about 1.5 to 2 h period after each single stimulation [3,8,9,10]. Further studies on longer experimental time to determine even longer period post stimulation effect would be clinically valuable.

Recent experimental study in decorticated rats have shown that repeated short periods of ano-genital afferents stimulation induced a prolonged inhibitory effect on micturition reflex well beyond the stimulation period [11]. Several clinical studies reported the beneficial effect of TNS and recommend as a alternative treatment for patients with refractory urinary tract dysfunction [6,12,13]. Inspired by these positive findings, the present experimental study was designed to determine whether TNS might induce a similar prolonged modulation of the micturition reflex as ano-genital afferents stimulation, and to provide experimental evidence supporting the clinical application of TNS for treatment of neurogenic urinary bladder dysfunction, especially OAB

## Materials and methods

### Animals and Surgical Preparation

The experiment was carried out on 15 adult female rats (Sprague-Dawley, 250-350 g). Under the anesthesia with methohexitone (Brietal, Eli Lilly Co., Indianapolis, Indiana, 70 mg./kg. intraperitoneally), the femoral vein and artery were cannulated to allow fluid injections and blood pressure recordings, a tracheotomy was performed for artificial respiration. Then the animals were decorticated following the reported procedures [14]. In brief, after craniectomy and dura excision, the fully decortication was carried out by gentle suction using a standard drawn glass pipette connected to a vacuum pump and sparing most of the diencephalon. The fenestra was then filled with hemostatic cotton or gelatin sponge soaked with thrombin to minimize bleeding in the entire experiment. This procedure rendered the animals unconscious so no further anaesthesia was required.

For recordings the animals were paralysed by a continuos i.v. infusion of pancurone bromide (0.3 mg/kg.h) and artificially ventilated with end-expiratory CO_2_ adjusted to about 3.5%. Care was taken to ascertain that the paralysed animals were adequately unconscious by regularly controlling that strong paw-pinches failed to induce changes in blood pressure and heart rate. Their body temperature was maintained at about 38°C by a feed-back controlled heating lamp.

The animals were sacrificed at the end of the experiment by an overdose of the anesthetics followed by severance of the heart. The experiments were approved by the animal research ethical committee of Guangzhou medical university in accordance with Chinese law.

### Cystometry and Tibial Nerve Stimulation

The experiment arrangement is shown in Figure 1A. Bilateral tibial nerve stimulation was performed by inserting two pairs of needle electrodes percutaneously close to the tibial nerves at the level of the medial malleolus with the cathode to anode (1 cm apart). A polyethylene catheter (8F) with a side hole close to the tip was inserted into the bladder through a slit in the proximal urethra and fixed by a 3-0 silk ligature around the catheterized urethra. The catheter was connected to an infusion pump and a pressure transducer for cystometry recordings.

**Fig 1.**
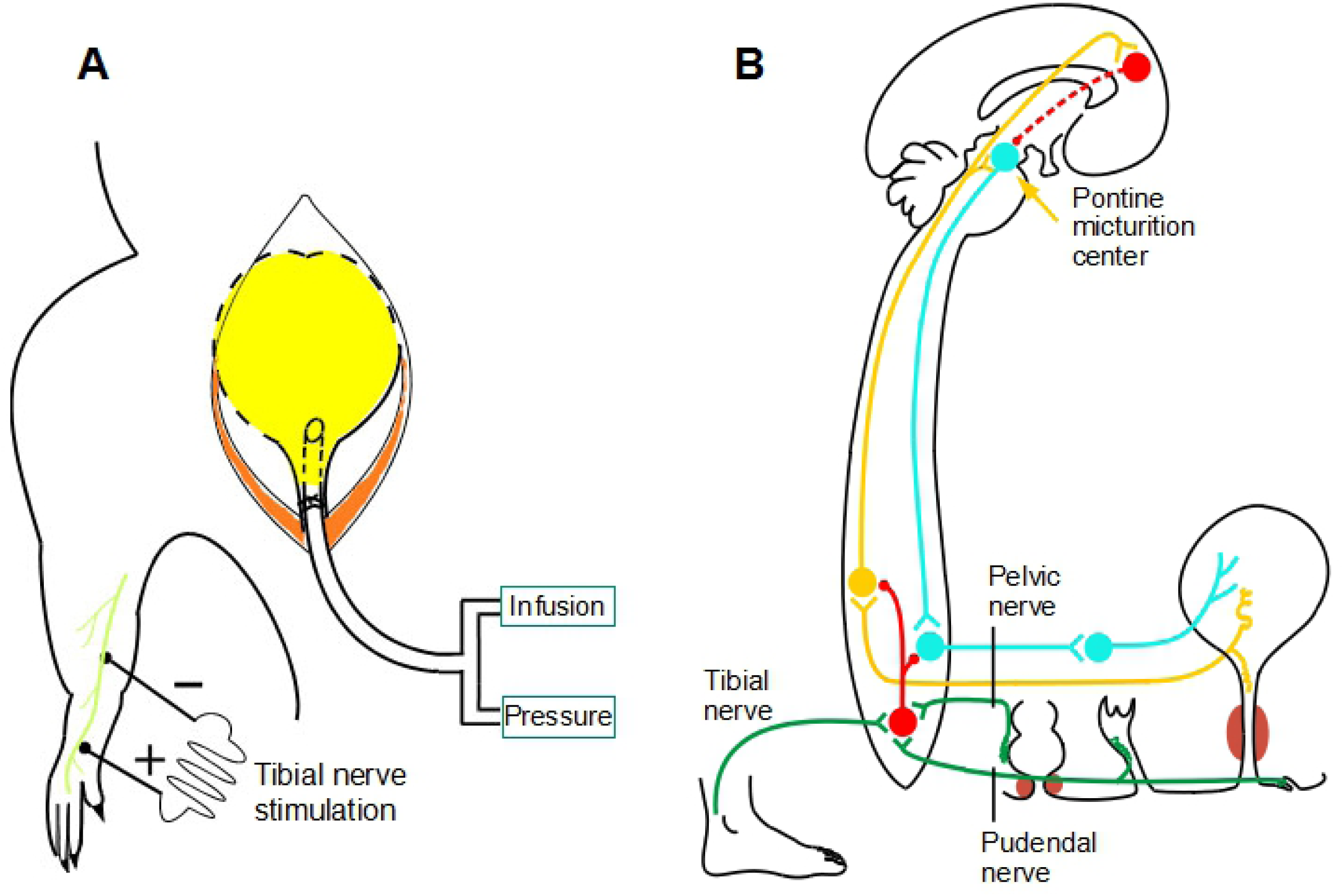
Schematic diagram of experimental arrangements in rat (A) and inhibitory control systems of the bladder in human (B). The afferents from tibial nerve, anal canal and vaginal mucosae are not only affecting the motor output from the spinal cord, but also send the signals up to the pontine center and to the cerebral cortex. All these pathways are there to prevent unintended micturition. (yellow – bladder afferents, blue – bladder efferent, green – inhibitory afferents from the indicated organs, red – inhibitory interneurons).

Constant flow cystometries were performed repeatedly with body-warm saline (0.07 ml/min) at about 10 min interval. As soon as bladder contracted, the infusion was turned off and the catheter opened. Care was taken to empty the bladder completely after each cystometry by lowering the catheter outlet below the level of the bladder. The micturition threshold volume (MTV) was defined as the amount of infused fluid into the bladder at onset of micturition contraction, which was used as the dependent variable. For control, 4 – 5 cystometroy recordings were performed before TNS. TNS was delivered for 5 min continuously, at frequency of 10 Hz, intensity of 3 times threshold for α-motor axons intensity determined by the induced obvious toe movement. These electrical stimulation parameters were chose since they are optimal as we confirmed previously and reported by others [11,15]. The same stimulation procedure were repeated for 6 times with 5 min interval including one cystometry between each simulation. During the stimulation, the bladder was empty with the catheter open. The volume of infused fluid into the bladder at onset of micturition contraction was defined as micturition threshold volume (MTV). The modulatory effect of TNS was defined as the mean of MTV compiled from several cystometries in each half hour before, during and after the TNS (see Figure 5).

### Statistical Analysis

For group comparison, the data are normalized and expressed as a percentage of the control using the Wilcoxon signed rank test. The data from different animals are presented as mean ± standard error and compared by repeated measures one-way ANOVA analyses. A *P* value of 0.05 or less was considered statistically significant. Statistical analyses were performed with SPSS 19.0 software (SPSS, Inc., Chicago, IL).

## Results

All the decorticated rats survived in a good state for 1-3 days with reasonably stable micturition reflexes. The stable bladder contractilities were ascertained by 4-5 sessions of control cystometries indicated by less than 10% of differences of MTV between each cystometry. The TNS induced a remarkable inhibitory effect on micturition reflex, manifested as a significant increase in MTV. Typical cystometry recodings were presented before and after TNS in Figure 2. One can see that a minimal increase in bladder pressure during the infusion until a micturition contraction occurred at the a bladder volume of 0.31 ml (Figure 2A) and 0.44 ml (Figure 2B). The recording **A** was obtained 10 min before, **B** was 5 min immediately after TNS. The stimulation induced 142% increase in MTV compared to the control in this case.

**Fig 2.**
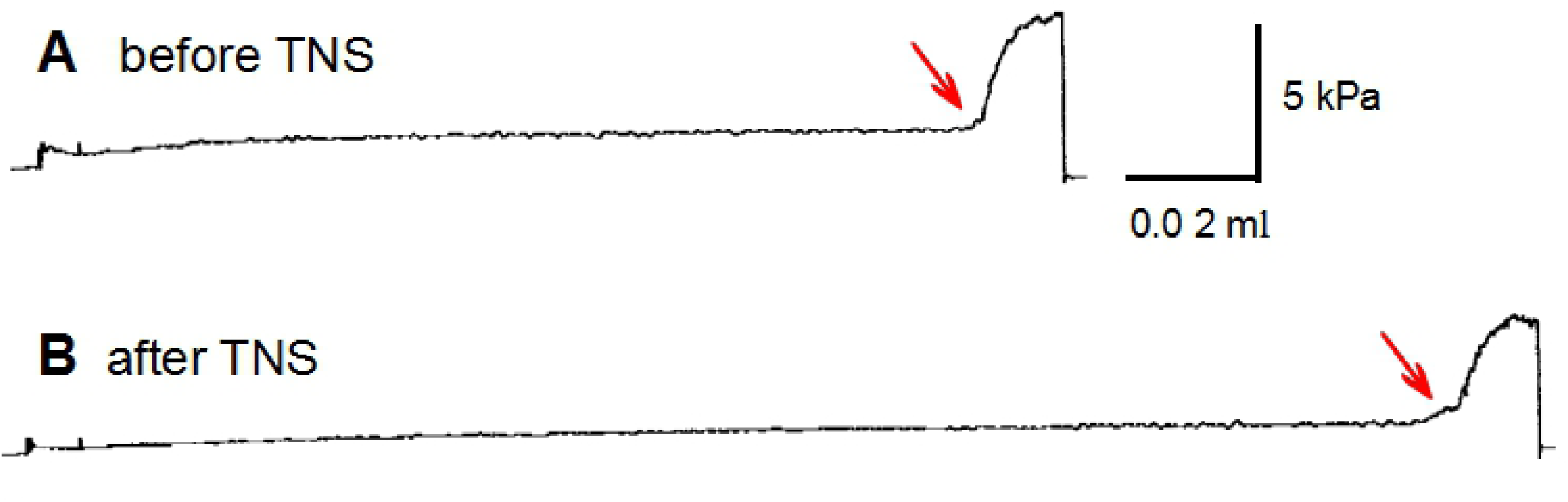
Cystometrograns (CMGs) recorded before (A) and after the tibial nerve stimulation (TNS) (B). The bladder was filled at a constant rate (0.07 ml/min) with body warm saline. The recording *A* was obtained 10 min before TNS, *B*, 5 min immediately after the TNS. TNS was delivered for 5 min at 10 Hz, at intensity of 3 times threshold for α-motor axons (same as other figures). Note that the increase in MTV induce by TNS after stimulation (0.44 vs 0.31 ml, 142%). Red arrow point the MTV (MTV, micturtion threshold volume, same as others).

The time curse of the changes in MTV induced by TNS is illustrated in Figure 3. Five cystometries had been tried to ascertain a stable bladder contraction before the stimulation. In this representative case, the MTV were between 0.1 – 0.15 ml, which is reasonable controls for the following test experiments. The inhibitory effect of TNS was demonstrated immediately after the first 5 min stimulation (e.g., the MTV increased from 0.15 to 0.27ml). This increase tendency was kept all the time in the next 4 repeated period stimulation, peaked to 0.38 ml after 5 periods of stimulation. After six periods of stimulation, the significant increase in MTV was obtained for more than 5 h.

**Fig 3.**
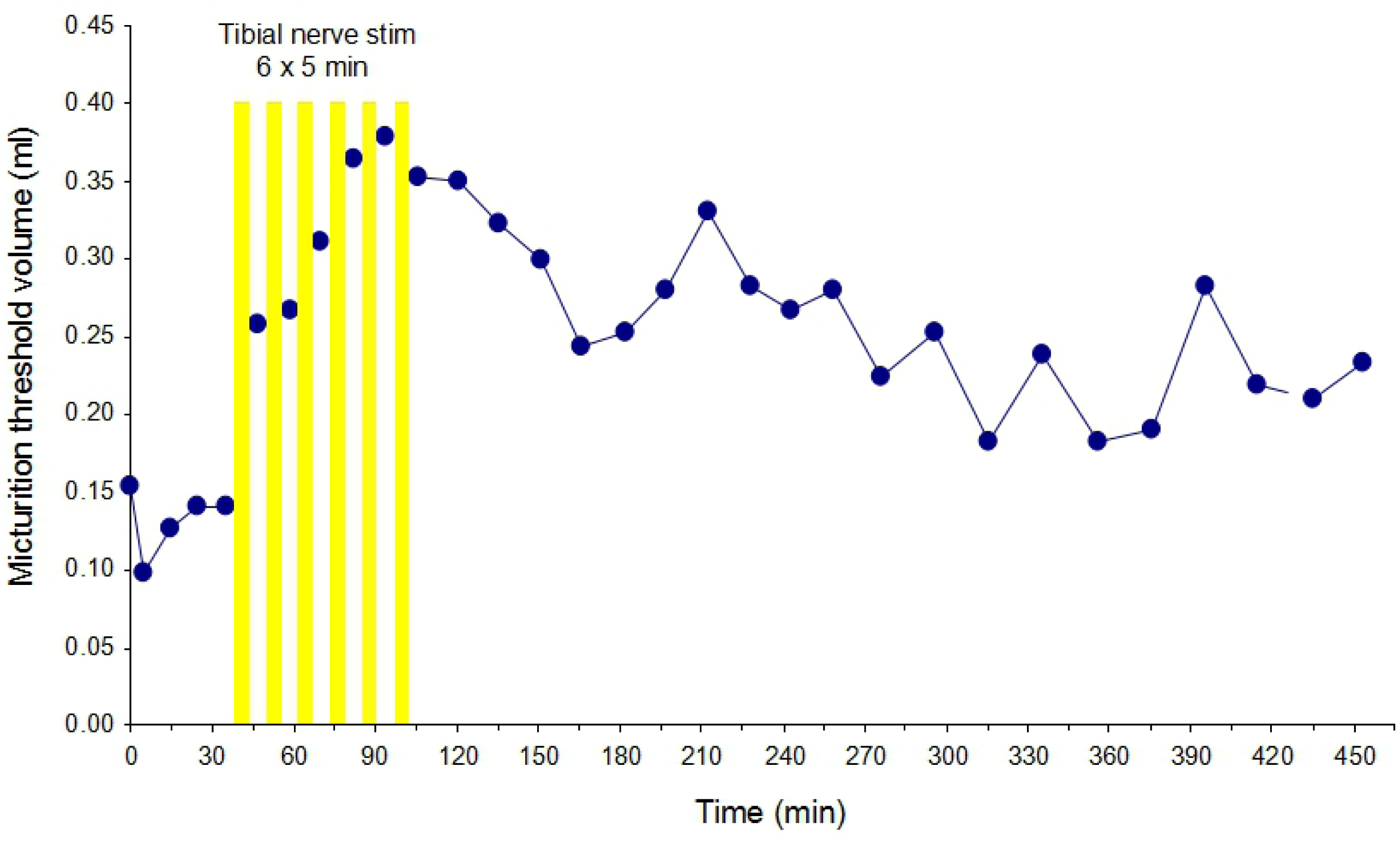
Changes in MTV before and after the tibial nerve stimulation. Each point in the plot represents the MTV of each cystometry at the indicated time interval. Each bar stands for 5 min stimulations.

A prolonged post stimulation inhibitory effect of TNS on the micturition reflex was demonostrated in another representative series of cystometrograns (CMGs) recordings (Figure 4). In this case, there was no significantly changed in MTV between 4 consecutive recordings before the stimulation (mean 0.39 ± 0.04 ml), while a significantly increased in MTV was observed inmmediatly after each 5 min stimulation and these 6 periods (half hour) repeated TNS induced a prolonged post stimulation inhibitory effect for more than 5 h. The normalized value were corresponded to pooling data histogram of figure 5.

**Fig 4.**
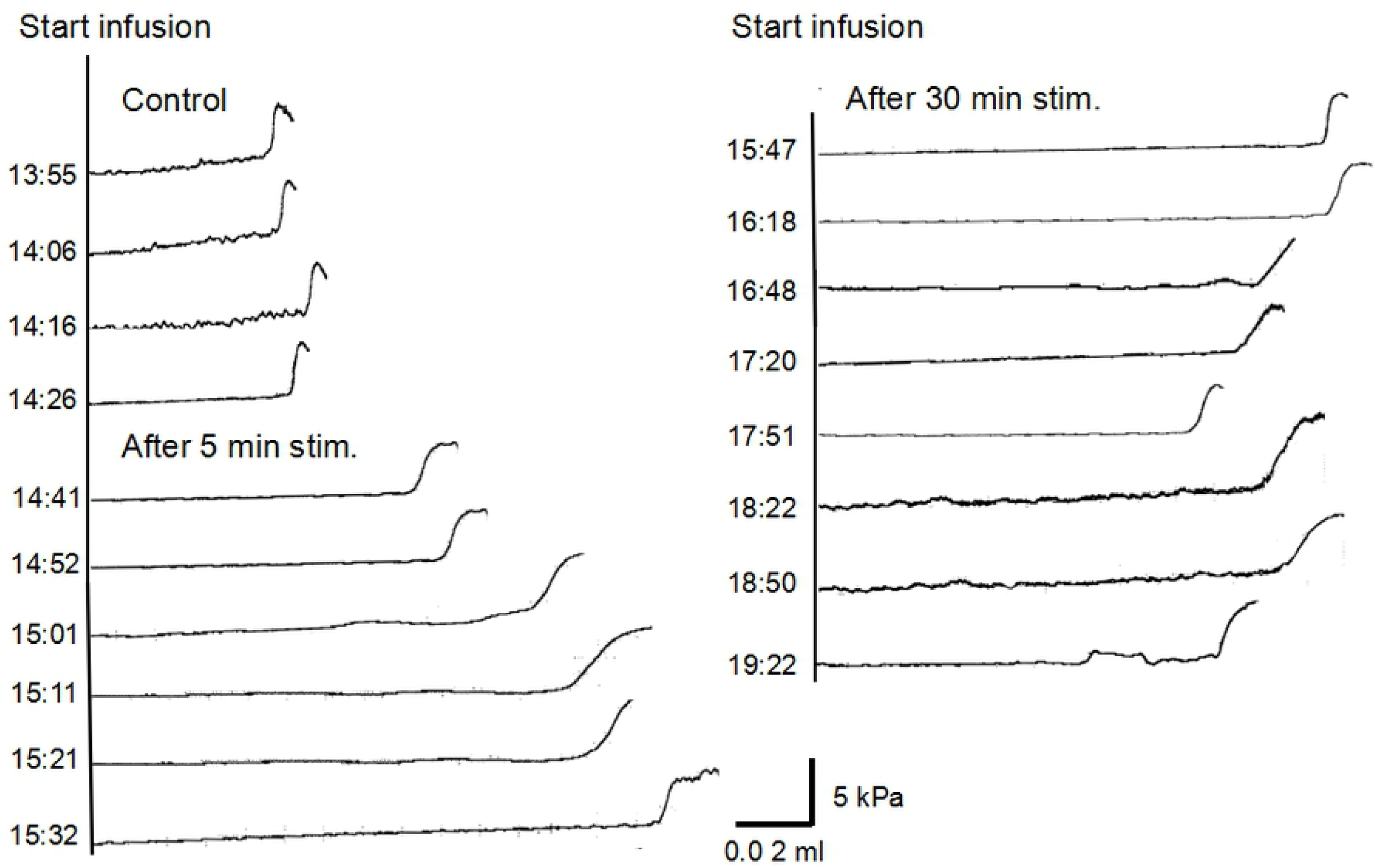
Representative a series of CMGs recordings in another animal (different from Figure 3) showing a prolonged poststimulation inhibitory effect of tibial nerv stimulation on the micturition reflex. MTV was not significantly changed between 4 consecutive CMGs as indicated time before the stimulation (mean 0.39 ± 0.04 ml). There was significantly increased in MTV after each 5 min stimulation and these 6 periods (half hour) tibial nerve stimulation induced a prolonged poststimulation inhibitory effect for more than 5 h. For simplifying, CMGs after repeated stimulations were omited every two consequtive CMGs. The normalized value were corresponded to pooling data plot as figure 5. Infusion rate: 0.07 ml/min. Bladder pressure and time calibration in the lower right corner refer to all CMGs traces.

**Fig 5.**
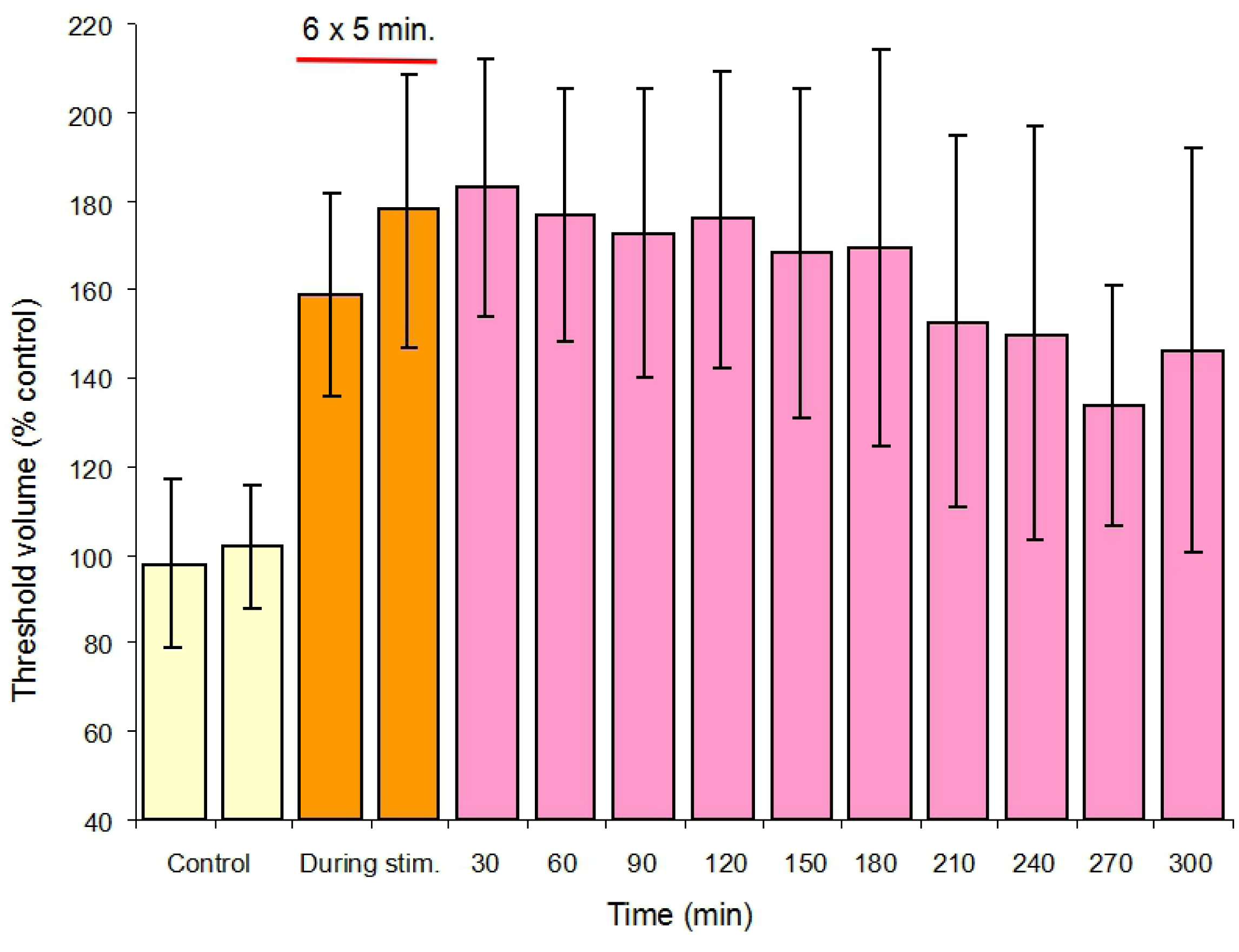
Time course of tibial nerve stimulation induced changes in MTV for all the animals. Each bar represents a normalized threshold value (±standard error of mean) in each half h before, during and after the stimulations. Six periods of TNS with 5 min interval were delivered as indicated by a horizontal red bar (n = 15; all *P* < 0.01).

For better comparison, the data of all animals (675 cystometries of 15 rats) were pooled and shows in Figure 5. To reduce scatter due to the large variations of MTVs between each individual, the values were normalized prior to pooling and expressed as percentage of the mean control value of each animal. In this summary figure, TNS induced a significant increase in micturition threshold volume for all animals during and after TNS (*P* < 0.01). The MTV increased to 160% of control after the first stimulation and it continuously increased to 180% of control after the second stimulation. After 6 period of repeated stimulations, the mean MTV reached peak of 190% control at the 30 min and it remained at about 150% 3 h after. Actually, one can see that from the entire histogram, a powerful inhibitory effect of TNS on the micturition reflex was kept almost no attenuation untill 5 h (Figure 5).

## Dicussion

As revealed in our previous study, a repeated short period of ano-genital afferent stimulation induced a prolonged poststimulation inhibition effect on micturition reflex for more than 5 h in the decorticated rats [11]. The same effect was demonstrated in the present study with a similar experimental paradigm but tibial nerve stimulation (TNS). The outcomes showed that TNS for 5 min at frequency of 10 Hz, intensity of 3 times threshold for α-motor axons induced 160% increase in MTV immediately after the first TNS compared to the controls, and the prolonged post stimulation inhibitory effect was observed after 6 period of repeated stimulation, a significant increase in MTV for more than 5 h (*P* < 0.001, Figure 5). This long last modulation effect of TNS is longer than those of other studies with similar stimulation parameter. For example, Tai and co-workers have found that a post stimulation of inhibitory effect on bladder activity was last for more than 2 h after TNS for 30 min, at 5 to 30 Hz in cats experiment [8]. In 2 of rats studies, the TNS induced post stimulation inhibition effect on micturition reflex for 50 min and 1 h, respectively [9,16]. All are much shorter than our observations.

The possible explanations for this difference might be that although the TNS parameters were roughly analogous, the patterns of stimulation were different, e.g. rather than continuous stimulation, we used an repeated short period stimulation, e.g. 5-min on and 5-min off stimulation pattern. This stimulation modality was applied since such an algorithm have been shown to effective for other central synapses [17]. In addition, it would be necessary to have a break time between each period stimulation for bladder to recover from the previous contraction. From the clinical point of view, a proper interval between each stimulation sessions would be important to activate the inhibitory effect on micturition reflex. Further more, it might be good for patient to adapt and bear the electrical stimulation, thus improve the treatment compliance [18]. In fact, this repeated stimulus modality has been applied clinically and beneficial effects have received in treatment of patients with OAB, although the stimulus session and recovery interval were proportionally longer compare to the rat, e.g, 30 min in each day or week [19-21]. In terms of TNS induced longer post simulation modulation effect, repeated short period stimulation seems better than the continuous stimulation.

It is well known that the micturition reflex is highly anaesthesia sensitive, especially to determine the long lasting poststimulation effect [22]. Therefore, different from other animal studies under the general anaesthesia, we used un-anaesthetised decorticated rat for the study. This rat model was not only overcome the influence of anaesthesia but also survived in a good condition with stable recordings for 1 – 3 days by properly paralysed, ventilation and intravenous infusions. As you can see in figure 3 and 4, four to 5 stable CMGs were obtained before the stimulation and a long last increase in MTV after TNS was highly significant even by intuition, suggesting that the acquired data in this study were more reliable and well comparable.

It is not surprised that the rats could survive for such a long time with a stable micturition reflex under repeated stimulation and cystometry recordings. Indeed, there was a report that the rats survived in a good condition for 6090 min (4 days) under the urethane anesthesia with the repeated stimulation and recordings [16]. Therefore, it is feasible to carry out a long time poststimlation modulation studies in the properly treated animals, although the decorticated rats as we presented here would be the best alternative, which is another explanation for us to be able to obtain the data of longer last TNS modulation effect than those of other studies in animal experiment.

Neuromodulation therapies utilize electrical stimulation to activate specific nerves that can modulate bladder activity in treatment of OAB and others voiding dysfunction. Tibial nerve stimulation (TNS) is one of the procedures due to its inhibitory effect on micturition reflexes. There are different protocols of TNS including direct ways, such as implanted stimulate device over the tibial nerve in human [23], bipolar cuff electrode and tripolar cuff electrode placed around the nerve in rat and cat, respectively [24]. The indirect ways are implantable wireless driver microstimulator [25], percutaneous tibial nerve stimulation (TNS) in a field way [4-6]. The TNS technique as we applied in the present study was similar to the clinical procedure (e.g., inserted 2 needles cranially 3-4 cm to the medial tibial malleolus and stimulated by connecting the electrostimulator). The reason we chose this technique for the study was based on the idea from Chinese acupuncture treatment by needles insertion onto the tibial nerve corresponded to point *Sanyinjiao* (Figure 1A), albeit the nerve dissection was made before the experiment (Figure 1). So far, there are no details in the publications on effect comparison among different tibial nerve stimulation approaches, although a doubtful different effect of TNS between surface electrode and nerve cuff procedures reported by Mario K [20]. Otherwise, all the procedures have similar inhibitory effect on the micturition reflex as long as a suitable stimulation parameters are applied. Those so called advanced techniques of direct nerve stimulation did not show more benefit of effect than those of indirect methods. The significantly increase in the MTV suggested that our TNS procedure in the rat is applicable.

Tibial afferent nerves project to the segments of the lumbosacral spinal cord that also receive inputs from bladder afferents travelling in the pelvic nerves [3]. Activate these systems by electrical stimulation produce central inhibition of the micturition reflex as we demonostrated in the present study, which is very similar to the effect of ano-genertal afferents stimulation. This inhibition occurs at least already at the spinal level, and rather than that, all these afferent systems are not only affecting the motor output from the spinal cord, but also send the signals up to the pontine center and cerebral cortex. All these pathways are well defined to involved in preventing unintended micturition (Figure 1B) [1,26].

Our stimulation intensity at 3 times threshold for α-motor axons (the threshold intensity for inducing a toe movement) was applied, since this TNS intensity is enough to activated both Aδ and C-fiber afferent as we determined by multiples threshold intensities for tibial nerve stimulation before the experiment. Other studies also revealed that this current had inhibit effect on isovolumetric bladder contractions in cat [3,25] and poststimulation inhibition effect indicated by increase in bladder capacity and compliance in rat [9]. We hypothesize that this somato-vesical inhibition effect was involved in both supraspinal and spinal circuits of the bladder via Aδ and C-fiber micturition reflex pathways, respectively.

Nevertheless, such a prolonged (5 h) post stimulation modulation effect of micturition reflexe would be better explained by modulation of the synaptic transmission in the central micturition reflex pathway. The idea was supported by our previous study of intravesical electrical stimulation induced enhancement of micturition reflex which could be prevented by systemic administration of a specific, competitive NMDA antagonist, CPPene [27]. Activation of NMDA receptors has been implicated to have a role in the modulation of synaptic efficacy. The mechanism of so called long term potentiation (LTP) is associated with the enhancement of excitatory synaptic transmission in the central nervous system. By analogy, the present bladder inhibitory effect of TNS might be due to a negative modulation of excitatory synapses in the central micturition reflex pathway. A prolonged decrease in synaptic efficacy of excitatory synapses, referred to as long term depression (LTD), has been observed in the hippocampus after intense activation of an inhibitory input to the target cells [27,28]. Other central processes, such as opioid receptors in the brain stem are also reported to involve in TNS inhibition of bladder overactivity in cats [29]. Since the effects were observed in animals lacking cortical control of the micturition reflex, the higher, cognitive functions would be not necessary. Thus, the re-education seen in some patients with OAB following repeated sessions of TNS may at least in part result from long term depression (LTD) like modulations of synaptic transmission in the subcortical micturition reflex pathway [28,30]. Of course, more investigations are crucial for further understanding the underlying mechanism of long lasting poststimulation effect induced by repeated or continuous TNS in animals and patients.

## Conclusions

A prolonged increase in MTV was demonstrated by repeated short periods of TNS in the decorticated rats. This inhibitory poststimulation modulation effect on micturition reflex was similar to that earlier reported for ano-genital afferent stimulation and resulted from activation of inhibitory neuromechanisms in the central micturition reflex pathway. We assume that the finding would offer a theoretical explanation for the clinical beneficial effect of TNS in patients with overactive bladder.

## Acknowledgments

This study was supported by Medical Scientific Research Foundation of Guangdong Province, China (grant number:WSTJJ20120544010197604063216). Natural Science Foundation of Guangdong Province, China (grant number: 2016A03030703). Swedish Medical Research Council (projects no 04767).

